# Flow cytometry-based isolation of *Salmonella*-containing phagosomes combined with ultra-sensitive proteomics reveals novel insights into host-pathogen interactions

**DOI:** 10.1101/2025.04.25.650444

**Authors:** Ritika Chatterjee, José Luis Marin Rubio, Francesca Romana Cianfanelli, Andrew Frey, Benjamin Bernard Armando Raymond, Abeer Dannoura, Camila Valenzuela, Meihan Meng, Tiaan Heunis, Mengchun Li, Frances Sidgwick, Jost Enninga, Andrew Filby, Matthias Trost

## Abstract

Macrophages engulf pathogens into dynamic phagosomes, which many bacteria manipulate for survival. However, isolating pure pathogen-containing phagosomes remains challenging. Here, we developed a novel flow cytometry-based isolation and ultrasensitive proteomics approach to analyse phagosomal and bacterial proteomes from macrophages infected with wild-type (WT) *Salmonella enterica* serovar Typhimurium (STM) or a Δ*phoP* mutant at 30 min and 4 hrs post-infection. Our approach provides higher throughput, requires lower cell numbers and quantifies more proteins than previous techniques. Our data reveals key host-pathogen interactions, showing induction of PhoP-dependent virulence factors and novel putative proteins that shape STM’s intracellular niche. Moreover, our data indicates that bacteria-containing phagosomes recruit mitochondrial membrane for production of reactive oxygen species. These findings provide new insights into *Salmonella*’s manipulation of phagosomal maturation and intracellular niche formation.

## Introduction

Phagocytosis is an evolutionary conserved cell function in which professional phagocytes such as macrophages, neutrophils, and dendritic cells engulf and remove particles like dead cells, foreign particles and invading pathogens.^1,2^ Upon binding to a wide range of receptors, pathogens are internalised by cytoskeleton rearrangements, leading to the formation of a *de novo* organelle, the phagosome. The phagosome undergoes a series of maturation steps, sequentially fusing with early and late endosomes and ultimately with lysosomes, forming the phagolysosome, within which the ingested particles are degraded. This degradation process generates antigens that are presented to the cell surface, thereby linking innate and adaptive immunity. However, numerous intracellular pathogens, including *Salmonella enterica* serovar Typhimurium (STM), have evolved mechanisms to manipulate phagosome maturation, altering its composition to establish an intracellular replicative niche within macrophages.

*Salmonella* infection can cause a spectrum of diseases, ranging from self-limiting gastroenteritis to systemic typhoidal fevers, affecting a wide array of hosts. After breaching the intestinal barrier, the bacteria disseminate to secondary infection sites through the lymphatic system. During this dissemination macrophages phagocytose the invading bacteria, however, *Salmonella* has evolved a two-component system PhoP/PhoQ^3^ which senses reduced Mg^2+^ concentrations, cationic antimicrobial peptides, and acidic pH within phagosomes. The PhoP system activates the expression of virulence genes, including proteins of the *Salmonella* pathogenicity island-2 (SPI-2) encoded type 3 secretion system which enables the injection of effector proteins into the host cytoplasm, modulating host trafficking pathways to support bacterial survival and replication^4^.

To comprehensively investigate global changes in phagosome composition, various methods have been developed to isolate pathogen-containing phagosomes for proteomic analysis. However, while latex or polystyrene bead phagosome isolation followed by proteomics has been a hugely successful technology to understand the molecular composition of phagosomes of inert beads^4–6^, isolating highly pure pathogen-containing vacuoles and phagosomes (from now on called phagosomes for simplicity) remains a technical challenge. This difficulty arises primarily due to the similar densities of mitochondria and bacterial phagosomes, as well as the complex tubular structures of certain pathogen-containing vacuoles, such as *Salmonella*-containing vacuoles (SCVs) at later timepoints, which hinder fractionation-based isolation.^5,6^ Nonetheless, since the early 2000s, researchers have successfully enriched bacteria-containing phagosomes for proteomics analysis using a variety of methods, including density gradient centrifugation, magnetic bead affinity purification, and biotin affinity purification^7–9^.

Here, we utilised an innovative antibody-free technique to isolate *Salmonella*-containing phagosomes from infected human THP-1 macrophages using flow-cytometry that we termed PhagoCyt. Utilising highly sensitive proteomic analyses, we achieve unprecedented depth in characterizing the proteomes of both the phagosome and intra-phagosomal *Salmonella*, revealing the intricate recruitment of host protein complexes and organellar membranes to STM-containing phagosomes.

## Results

### Isolation of bacteria-containing phagosomes using antibody-free fluorescence-activated cell sorting (FACS)

We have developed a robust flow cytometry-based method, for isolating pathogen-containing phagosomes, which we termed PhagoCyt. This powerful approach enables the isolation of highly pure phagosome populations, which can be analysed using ultrasensitive proteomic techniques to characterise the proteomes of both phagosomes and intraphagosomal pathogens (**Fig.1A**).

**Figure 1:**
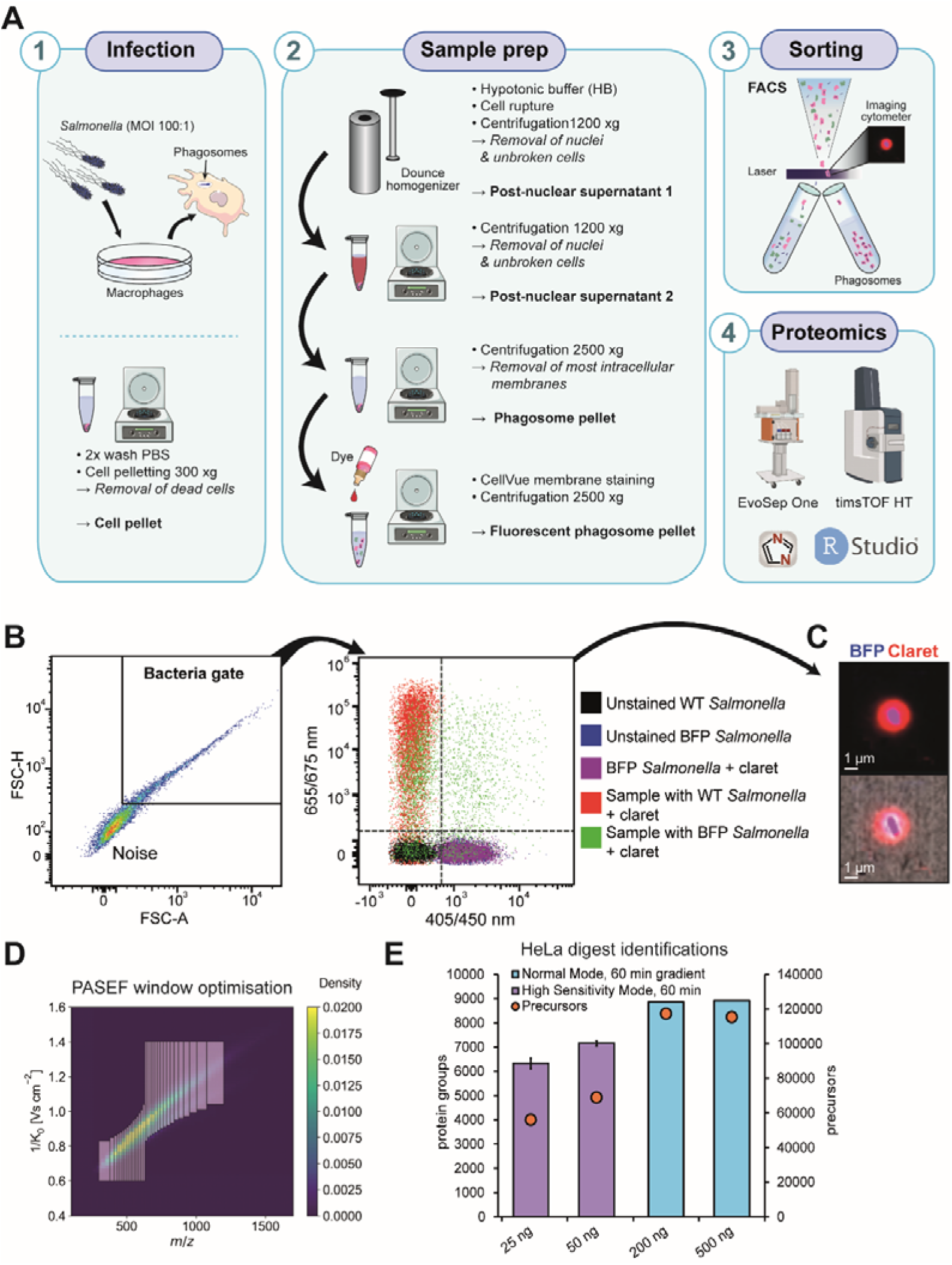
PhagoCyt workflow and mass spectrometry optimisation. (**A**) PhagoCyt workflow to isolate *Salmonella*-containing phagosomes by flow cytometry. **(B)** Flow cytometry strategy and control experiments. Bacterial phagosomes were first selected sorted based on forward scatter height (FSC-H) and forward scatter area (FSC-A), and then only double positives for BFP (bacteria) and Cellvue Claret Far Red were selected for sorting. Unstained bacteria (black), BFP-tagged *Salmonella* alone (blue), BFP-tagged *Salmonella* treated with CellVue Claret Far Red (purple) and cell lysates containing phagosomes with *Salmonella* not expressing BFP (red) do not generate any double positive hits. Only BFP-expressing *Salmonella* within phagosomes are double positive (green). **(C)** Imaging flow cytometry picture of a selected phagosome using ImageStream X Mark II. **(D)** Mass spectrometry parameters and specifically PASEF windows were optimised for the timsTOF HT. **(E)** Protein identifications on an EvoSep One/timsTOF HT platform for HeLa digests using these optimised parameters.

Microbes can be labelled either by expressing fluorescent proteins or using live-cell stains such as CellTrace Violet (CTV), allowing for the study of non-genetically modified clinical isolates (**Suppl. Fig.1A**). Importantly, CTV staining does not affect phagocytic uptake (**Suppl. Fig 1B**). We optimised the gating strategy for bacterial phagosomes first on size (**Fig. 1B; Suppl. Fig 1D**). To visualise phagosomal membranes in macrophage lysates, we identified 4 µM CellVue Claret Far Red as optimal, enabling the exclusion of bacteria merely attached to membrane fragments (**Suppl. Fig. 1C**). Importantly, CellVue does not stain bacterial membranes, ensuring specific labelling of host-derived phagosomal structures (**Fig. 1B; Suppl. Fig 1E**).

Human THP-1 cells were differentiated into macrophages using 10 ng/ml Phorbol-12-myristate-13-acetate (PMA), resulting in high expression levels of macrophage markers CD11b, CD11c and CD14 (**Suppl. Fig 2A-E**). These macrophages were then infected with BFP-expressing *Salmonella* Typhimurium at a multiplicity of infection (MOI) of 1:100 for 30 min and the infected cells were maintained in gentamicin-containing media for up to 4 hrs

BFP-expression in STM does not affect bacterial growth and survival (**Suppl. Fig 1F,G**). As STM infection induces macrophage cell death at later time-points (past the 4 hours used here)^10^, centrifugation steps were included to ensure that only live macrophages were pelleted.

Live, infected macrophages were mechanically lysed using a dounce homogenizer, and unbroken cells and nuclei were removed by centrifugation. Phagosomes were then pelleted by centrifugation from this post-nuclear supernatant, membranes stained with CellVue and analysed by flow cytometry. This revealed double-positive phagosomes containing bacteria and CellVue-stained membranes. Control experiments (**Fig. 1B**) showed that unstained bacteria (black), BFP-tagged *Salmonella* alone (blue), BFP-tagged *Salmonella* treated with CellVue Claret Far Red (purple) and cell lysates containing phagosomes with *Salmonella* not expressing BFP (red) do not generate any double positive signals. Only BFP-expressing *Salmonella* within phagosomes exhibited a double positive signal (green). Imaging cytometry confirmed >95% purity and intact phagosomal membranes (**Fig. 1C**).

Assuming that a single phagosome contains about 0.5% of a cell’s content (∼0.25 pg of protein), 10^5^ phagosomes contain an estimated 25 ng of protein. We optimised sample preparation for these low-input samples, employing the high-recovery S-Trap method for cell lysis and protein digestion. To enhance proteomics sensitivity, we also optimised diaPASEF parameters on a timsTOF HT instrument (**Fig. 1D**). Specifically, we designed a number of variable-width diaPASEF acquisition schemes using pydiAID^11^ and tested them with optimal and low loads of tryptic peptides. Testing 500 ng of the protein digests with a 60-min gradient yielded >8000 protein group IDs and >120,000 precursors with the optimal 16 window (W) diaPASEF acquisition scheme. In high-sensitivity mode, the timsTOF HT identified 6000–7000 protein groups and ∼60,000 precursors from as little as 25–50 ng of HeLa lysate digest (**Fig. 1E**).

Using this workflow, we performed a proteomics analysis of phagosomes isolated from THP-1 macrophages that were infected with BFP-expressing wild-type (WT) *Salmonella* Typhimurium (STM) and a Δ*phoP* mutant, which is unable to respond to the hostile phagosomal environment^3^. For each condition, macrophages were challenged with an MOI of 1:100 for 30 min, followed by gentamicin treatment to eliminate extracellular bacteria. Macrophages were then chased for 30 min or 4 h to generate phagosome samples at early/medium and late/matured timepoints.

### Maturation of WT STM phagosomes shows intricate interactions with organelles and protein complexes

Phagosome maturation involves dynamic interaction with host organelles and protein complexes^1^. Across six biological replicates, PhagoCyt showed very high reproducibility despite being isolated on different days (**Suppl. Fig 3A-C**) and clear separation of individual conditions (**Suppl. Fig. 3D**). Overall, we identified 5800 proteins, of which were 4625 human and 1175 STM proteins (<1% FDR, minimum of 2 peptides per protein). Stringent filtering, i.e. only taking proteins that have been quantified in 4/6 replicates, reduced that to 4761 quantified proteins of which 3900 were human and 861 bacterial (**Fig. 2A, Suppl. Tables 1-5**). A total of 1331 proteins significantly changed in abundance during phagosome maturation in STM WT phagosomes (q < 0.05, fold-change > 1.5), with 858 human and 444 *Salmonella* proteins increasing, while 557 human and 15 *Salmonella* proteins were decreasing (**Fig. 2A**).

**Figure 2:**
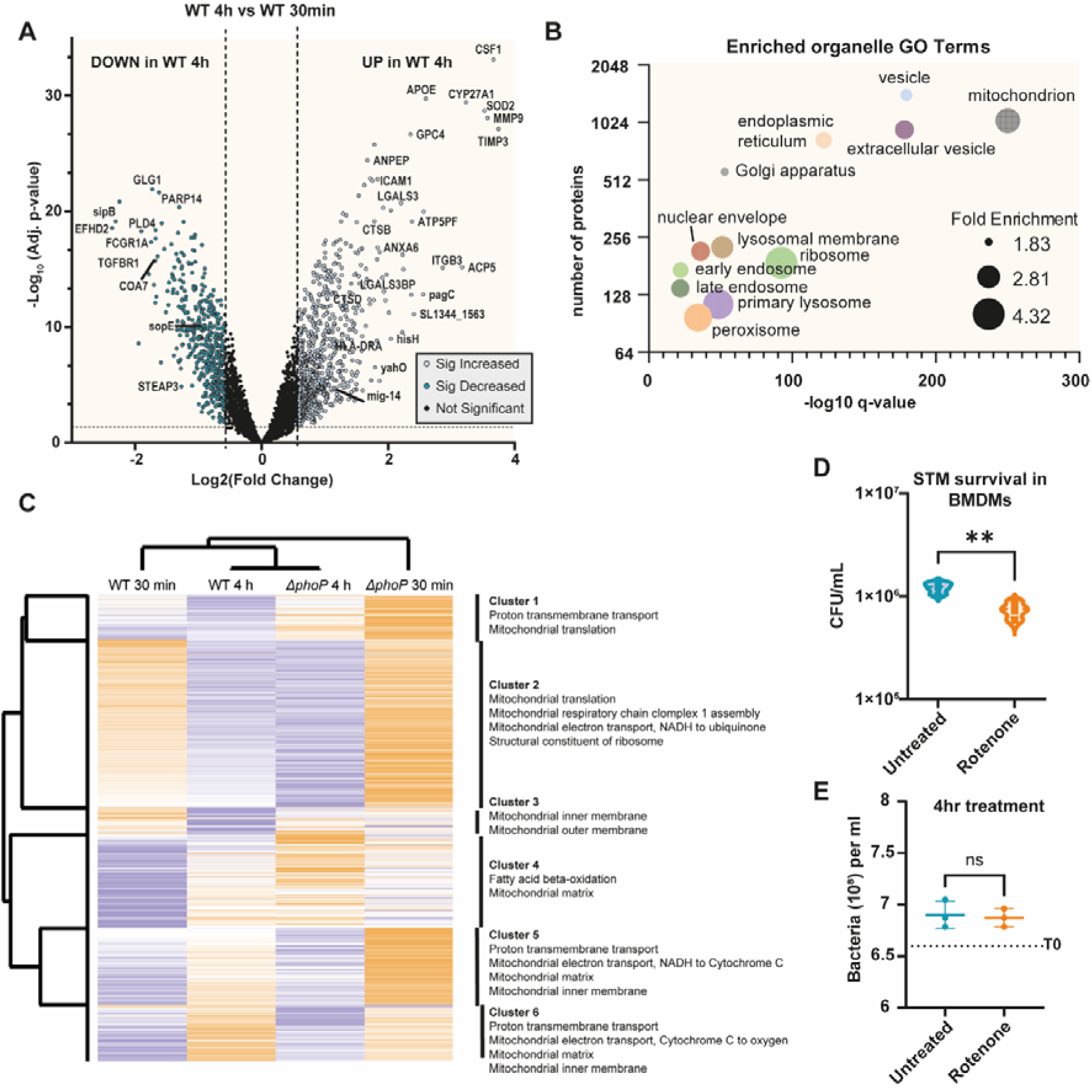
Proteomics results show intricate interaction of phagosomes with host organelles. (**A**) Volcano plot of the proteomics data of STM WT 4h vs 30 min. **(B)** GO-term analysis of organelles identified in STM phagosomes. **(C)** Heatmap of identified mitochondrial proteins shows significant changes of mitochondrial proteins over time and within bacterial strains tested. **(D)** Treatment of STM-infected macrophages for 4 hrs with 2 µM of the mitochondrial complex I inhibitor rotenone, reduces STM survival, probably due to increased ROS production. **(E)** Rotenone has no effect on bacteria in LB broth (4 hrs growth). **= p<0.01 (Welch Test).

Phagosomal proteomes were highly enriched for vesicular, endosomal and lysosomal proteins, along with proteins from the endoplasmic reticulum, mitochondria and nuclear envelope proteins (**Fig. 2B**). ER proteins and some mitochondrial proteins have also been identified on immunologically silent polystyrene-bead phagosomes^12^. However, while recruitment of ER membrane is known for STM phagosomes^11^, the identification of mitochondrial proteins in bacteria-containing phagosomes was in previous publications considered a contaminant, specifically when isolated by ultracentrifugation^9^, due to the similar density of mitochondria and bacteria. However, our FACS-based isolation does not rely on density gradients. Furthermore, mitochondrial protein abundances varied between conditions (**Fig. 2C**), suggesting that mitochondria may be actively recruited to phagosomes rather than present as passive contaminants.

STM secretes several effector proteins that modify mitochondria^13,14^. As mitochondrial reactive oxygen species (ROS) have been shown to contribute to pathogen killing in phagosomes^15^ and mitochondrial ROS-producing membranes have been shown to be delivered to bacteria-containing phagosomes via mitochondria-derived vesicles (MDVs)^16^, we tested if inhibition of mitochondrial Complex I, which increases ROS production, had an effect on *Salmonella* survival or replication. Indeed, inhibition of Complex I with rotenone increased *Salmonella* killing significantly (**Fig. 2D**), while rotenone treatment of *Salmonella* outside of cells did not affect their viability or growth within the same time-frame (**Fig. 2E**). Overall, this indicates that recruitment of mitochondrial membranes to the phagosome represents a host defence mechanism.

### The STM phagosome proteome changes drastically during maturation

Comparison of the 30 min and 4 hr STM WT phagosomes showed that several proteins were highly enriched in 4-hour STM phagosomes, including matrix metalloproteinase-9 (MMP9)^17^, which plays a key role in extracellular matrix remodelling, and during bacterial infections it recruits leukocytes and facilitates cytokine, chemokine and defensin activation^18^. Interestingly its inhibitor, metalloproteinase inhibitor 3 (TIMP3)^19^ is equally more abundant. Other strongly upregulated proteins included IFI30/GILT, a lysosomal thiol reductase involved in disulfide bond reduction; Galectin-3 (LGALS3), which is crucial for phagocytosis and phagosomal membrane repair^20,21^; and Gasdermin-A, a pore-forming protein, that can induce pyroptosis by preferentially targeting mitochondrial membranes^22^. Additionally, alpha-L-iduronidase IDUA which degrades glycosaminoglycans, was upregulated nearly fourfold, raising the possibility that it may contribute to the sugar degradation of the bacterial surface (**Fig. 3A**).

**Figure 3:**
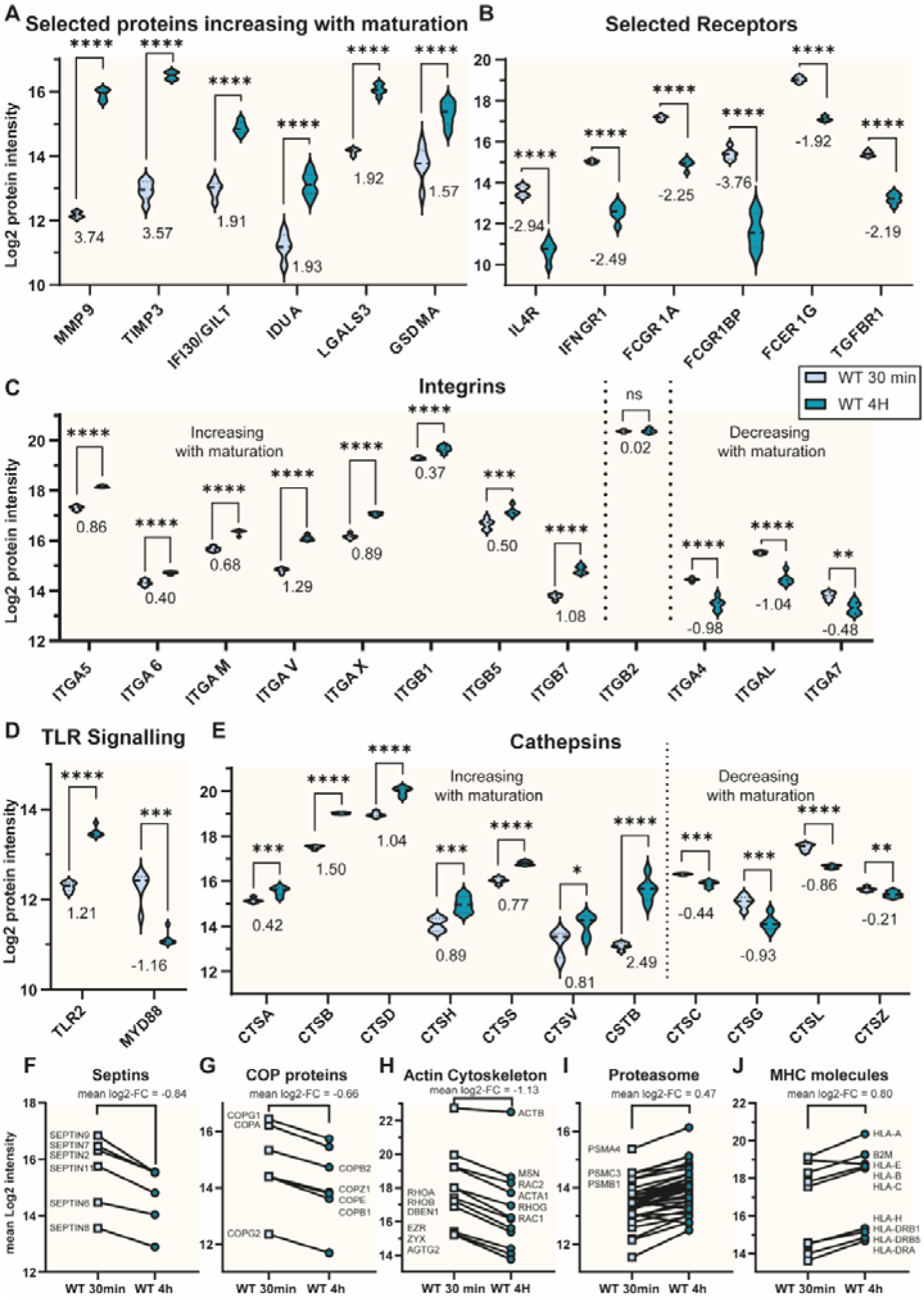
Selection of significantly changing proteins and protein complexes in WT STM phagosomes. (**A-E**) Normalised protein log2 intensities of selected proteins in WT 30 min (early, light blue) and WT 4 hr (late, dark blue) STM phagosomes. **(F-J)** Mean normalised protein log2 intensities of protein complex members of selected protein complexes. Mean log2 fold-changes (FC) were calculated from the log2 differences of the individual proteins. Numbers in the figure represent log2 fold-changes.*: p<0.05; **: p<0.01; ***: p<0.001; ****: p<0.0001, unpaired t-test (FDR controlled).

As expected, most plasma membrane receptors such as the Fc- and cytokine receptors significantly decreased with maturation, either through recycling or degradation (**Fig. 3B**). Interestingly, integrins did not follow that trend as some members of the integrin-family decreased in abundance, while others increased over time (**Fig. 3C**). Notably, ITGAL and ITGB2, which are critical for *Salmonella* clearance^23,24^, were among those increasing. Given their role in actin cytoskeleton regulation, integrin recruitment may contribute to regulating the highly abundant actin cytoskeleton machinery around STM phagosomes (**Fig. 3H**). A particularly striking observation was the significant increase in Toll-like receptor 2 (TLR2) over time (**Fig. 3D**), which contrasts with the behaviour of TLR2 on normally maturing bead phagosomes^25^. This suggests that STM phagosomes may engage distinct signalling pathways during maturation.

In a simplified model of phagosome maturation, lysosomal proteins such as cathepsins, increase with maturation. Indeed, these key lysosomal proteases mostly increase with maturation, but similarly to integrins, not all members behave as expected, with four cathepsins reducing with time (**Fig. 3E**), suggesting that trafficking to the STM phagosome from cellular cathepsin reservoirs is complex and potentially remodelled by STM.

Several protein complexes exhibited decreasing abundance with maturation, including septins that are essential for phagosome formation but have also roles in recognising bacterial cell division within phagosomes^26,27^ (**Fig. 3F**), the coatomer complex (COP proteins) that transports proteins from the Golgi (**Fig. 3G**) and the aforementioned actin cytoskeleton (**Fig. 3H**). Conversely, protein complexes that increased during maturation included the proteasome (**Fig. 3J**), which has recently been shown to generate anti-bacterial peptides^28^ as well as peptides for antigen presentation. Fittingly, MHC molecules also increase during maturation (**Fig. 3K**). These findings highlight the intricate regulation of phagosome maturation, revealing both expected and unexpected protein dynamics that shape the phagosomal environment during *Salmonella* infection.

### *Salmonella* actively modulates phagosomal protein recruitment in a PhoP-dependent manner

*Salmonella* Typhimurium actively modifies its phagosomal environment by secreting effector proteins, primarily through the *Salmonella* Pathogenicity Islands I and II (SPI-1/SPI-2)^4^. The ability to recognise the acidic phagosome environment through PhoP is essential for the induction of SPI-2 T3SS and secreted effector proteins, responsible for intracellular survival and replication^3^.

Reactive oxygen species (ROS) are essential in killing intracellular pathogens. The cell counteracts oxidative stress using superoxide dismutases which convert superoxide into H_2_O_2_. Interestingly, the cytoplasmic superoxide dismutase 1 (SOD1) does not change in the measured conditions. However, the mitochondrial SOD2 is not only significantly more abundant on the phagosome, but also increased hugely with maturation, which is an indication of very high ROS levels at the maturing phagosome (**Fig. 4A, Suppl. Table 7**). This is also supported by the induction of bacterial superoxide dismutases such as SodCA (**Fig. 5B**). Interestingly, while the NADPH oxidase (NOX) complex decreased in WT STM phagosomes, mitochondrial ROS production may compensate for this decline, specifically as interferon-induced nitric oxide synthases (iNOS) were undetectable at the analysed time points (**Fig. 4A**).

**Figure 4:**
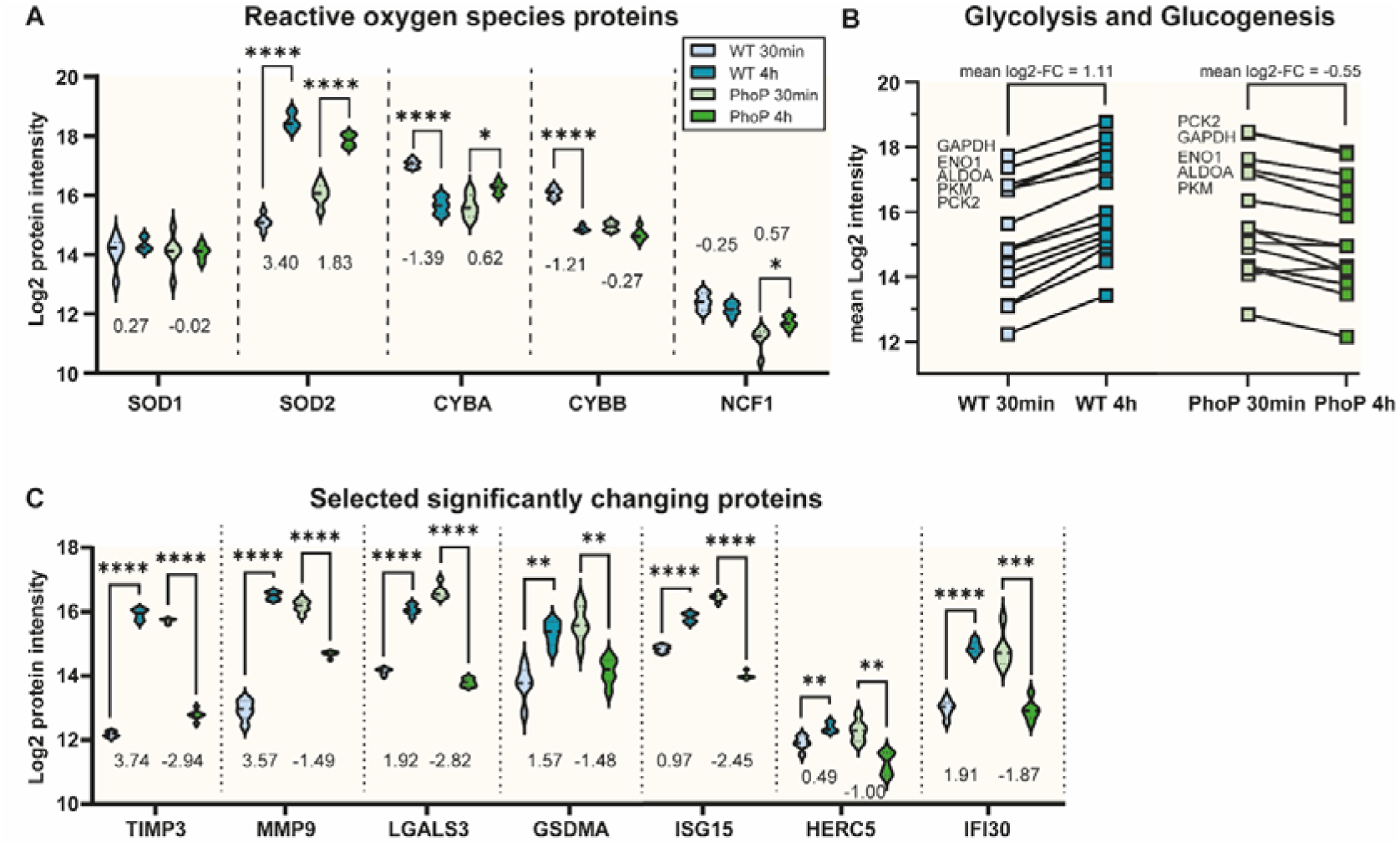
Selected protein changes in phagosomes of the PhoP-mutant and dead WT STM. **(A)** Normalised protein log2 intensities of selected proteins in superoxide conversion and reactive oxygen species generation in early (30 min) and late (4 h) phagosomes of (live) WT STM and Δ*phoP* mutant STM. **(B)** Mean normalised protein log2 intensities of proteins involved in glycolysis and glucogenesis. Mean log2 fold-changes (FC) were calculated from the log2 differences of the individual proteins. **(C)** Normalised protein log2 intensities of selected significantly changing proteins. Numbers in the figure represent log2 fold-changes. *: p<0.05; **: p<0.01; ***: p<0.001; ****: p<0.0001, unpaired t-test (FDR controlled).

**Figure 5:**
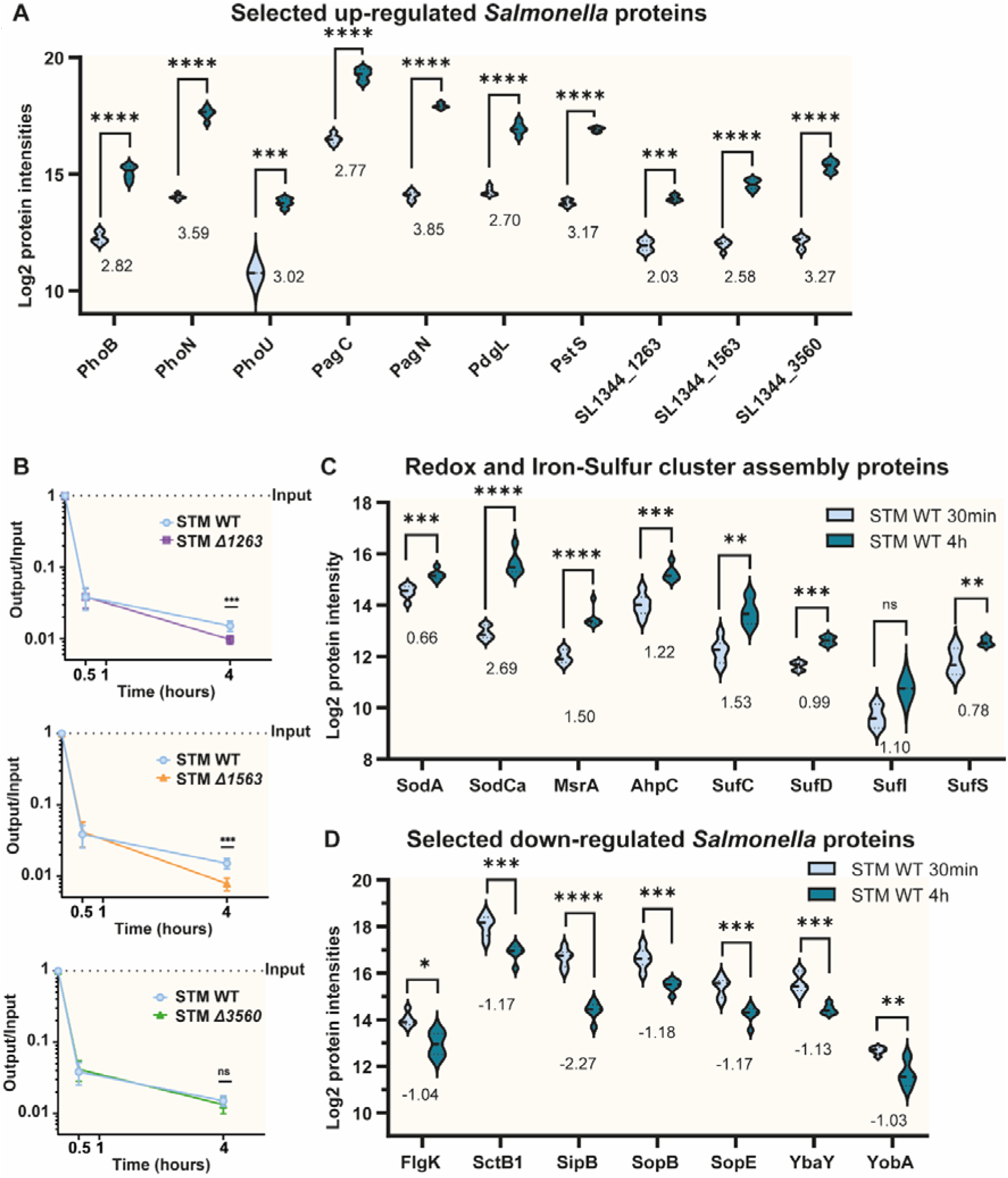
Selected bacterial protein changes in WT STM within phagosomes. **(A)** Normalised protein log2 intensities of selected bacterial proteins induced in late (4 hr) *Salmonella* phagosomes. **(B)** Infection experiments in BMDMs of STM WT and knock-out strains of the three novel bacterial proteins involved in infection. (**C**) Normalised protein log2 intensities of selected bacterial proteins of redox and iron-sulfur cluster assembly proteins. **(D)** Normalised protein log2 intensities of selected bacterial proteins reduced in abundance in late (4 hr) *Salmonella* phagosomes Numbers in the figure represent log2 fold-changes. *: p<0.05; **: p<0.01; ***: p<0.001; ****: p<0.0001, unpaired t-test (FDR controlled).

Glycolysis is essential for macrophages to clear *Salmonella* Typhimurium. Despite previous reports suggesting that STM reduces overall glycolytic activity in infected macrophages^29^, we observed an increased association of glycolytic proteins with maturing phagosomes. However, this pattern was reversed in the PhoP mutant, suggesting that PhoP regulates metabolic rewiring at the phagosome (**Fig. 4B**).

Notably, several proteins that increase with maturation in STM WT phagosomes - including MMP9, TIMP3, Galectin-3, and Gasdermin-A - exhibited opposite trends in the PhoP mutant (**Fig. 4C**). A similar reversal was observed for interferon-induced proteins, such as ISG15, its ligase HERC5, and IFI30/GILT, which decreased with maturation in PhoP mutant phagosomes. This suggests that PhoP-dependent effectors actively remodel the phagosomal environment, influencing key immune and metabolic processes.

### PhagoCyt identifies novel putative *Salmonella* effector proteins

STM utilises the PhoP/Q system to sense vacuolar low pH, which induces the expression of effector proteins secreted through the SPI-2 T3SS. These effectors are critical for bacterial survival and replication within the host^30^.

Consistent with previous studies^3^, we observed a strong induction of PhoP-regulated genes in STM phagosomes at the 4-hour time point, compared to 30-minute phagosomes (**Fig. 5A**). This included key PhoP/Q system components (PhoP, PhoQ, PhoN, PhoU, PhoB) as well as the phosphate transporter PstS, part of the ABC transporter complex PstSACB, which is activated by PhoB in response to phosphate starvation^31^. Additionally, we observed a significant increase in the outer membrane adhesin and invasin protein PagN, which facilitates adaptation to intracellular host niches^32^, as well as the PhoP-dependent upregulation of PagC, which has been implicated in bacterial outer membrane vesicle (OMV) formation^33,34^. Finally, we detected a marked increase in PdgL (PcgL), a periplasmic D-Ala-D-Ala dipeptidase. The identification of these well-documented proteins supporting STM infection validates our approach.

Beyond known effectors, PhagoCyt identified several putative proteins involved in infection. Among the most highly upregulated proteins at 4 hours were SL1344_1263 (lipid A deacylase LpxR family protein)^35^ which is regulated by HilD^36^, SL1344_1563, a predicted-cell envelope protein, and SL1344_3560, a predicted phosphatidic acid phosphatase (**Fig. 5A**). While the latter two have been shown to be supportive of establishing infection in the intestinal milieu in a transposon screen^37^, our own infection experiments in BMDMs showed that deletion of SL1344_1563 or SL1344_1263 resulted in a significant reduction in intracellular survival compared with the WT strain. Loss of SL1344_3560, however, did not have a significant effect (**Fig. 5B**).

To counter high levels of ROS generated by the host cell, STM significantly induces the abundance of the ROS-scavenging proteins, including the superoxide dismutases SodCa and SodA, the methionine sulfoxide reductase A (MsrA), and the alkyl hydroperoxide reductase AhpC^38^. We also observed increased levels of the Iron-Sulfur (Fe-S) cluster assembly proteins (SufC, SufD, SufI, and SufS) which support survival in macrophages^39^ and act as potent negative regulators of SPI-1 thereby helping STM establish a competent niche inside host cells^40^ (**Fig. 5C**).

While a large number of bacterial proteins increased in abundance during phagosome maturation, only a smaller subset showed decreased levels. These include FlgK, a flagellar gene which is downregulated upon entering the cell to evade the host innate-immune pathways of recognition of flagellar components as PAMPs^41^. Furthermore, we observed a reduction in the SPI-1 injectisome complex proteins such as SctB1^42^ and also effectors delivered by SPI-1 such as SipB, SopB and SopE, validating the existing literature. However, we also identified novel potential effectors of early infection, including the mostly uncharacterised Copper resistance protein YobA and lipoprotein YbaY, which is involved in biofilm formation, both of which are about 2-fold more abundant in the early phagosomes (**Fig. 5D**). This data suggests that PhagoCyt can serve as a powerful tool to identify novel microbial effector proteins of intracellular pathogens.

## Discussion

Phagosomes are highly dynamic organelles that interact with practically all cellular membrane trafficking pathways^1^. Due to this dynamic nature, proteomics of isolated phagosomes has been a powerful tool to identify unbiasedly novel biological insights. Polystyrene bead phagosome can be isolated to high purity by floatation in ultracentrifugation gradients and have therefore been extensively used to study these membrane dynamics and how macrophage activation affects phagosomal maturation^12,25,43,44^. The isolation of pathogen-containing phagosomes, however, has been much more difficult as many intracellular pathogens, and specifically bacteria, have a similar density to some host organelles and are therefore difficult to isolate through centrifugation alone. Additional steps such as magnetic bead affinity purification or biotin affinity purification^7–9,45^ were used in attempts to overcome these problems. Nonetheless, nuclear and mitochondrial proteins were often considered “contaminants” as their presence in phagosomes was considered a side effect of the isolation procedures. The here presented approach was carefully optimised to obtain a phagosome-enriched cellular supernatant from which phagosomes were then isolated using a highly sensitive flow cytometry approach. Our approach is not only relatively fast compared to other approaches but – when combined with low-input proteomics – also requires fewer cells. We only injected the digests of the equivalent of 100,000 phagosomes (∼25 ng) onto the mass spectrometer.

While flow cytometry (phagoFACS) has been used to analyse phagosomal subpopulations^46,47^ and isolate intracellular bacteria^48^, this is the first time where flow cytometry has been used to isolate intact phagosomes. The power of this approach is that not only the phagosomal host proteome, but also the bacterial proteome is available through the proteomics analysis. This data showed the induction of many known SPI-2 effector proteins, validating our approach. Additionally, it identified several novel putative effectors that will require further study. Furthermore, this study is the first to isolate *Salmonella*-containing vacuoles (SCV)/phagosomes from macrophages as previous studies were performed in HeLa cells^49^ and *Dictyostelium discoideum*^50^.

Overall, this method will enhance the ability to isolate pathogen containing phagosome and allow the unbiased molecular characterisation of host-pathogen interaction at this important organelle.

## Online Methods

### Antibodies

- BUV737 Rat Anti-CD11b (#612801, BD Biosciences).
- BUV737 Rat IgG2b, κ Isotype Control (#612762, BD Biosciences).
- CD11c Monoclonal Antibody (N418), APC (#17-0114-82, Invitrogen).
- Armenian Hamster IgG Isotype Control (eBio299Arm), APC (#17-4888-82, Invitrogen).
- CD14 Monoclonal Antibody (61D3), Alexa Fluor 700 (#56-0149-42, Invitrogen).
- Mouse IgG1 kappa Isotype Control (P3.6.2.8.1), Alexa Fluor 700 (#56-4714-80, Invitrogen).
- FITC anti-human CD44 Antibody (#338803, BioLegend).
- FITC Mouse IgG1, κ Isotype Control Antibody (#400107, BioLegend).
- Alexa Fluor 488 (AF488) Goat anti-Rabbit IgG antibody (Invitrogen).

### THP-1 cell culture and differentiation into macrophages

The human monocytic leukaemia cell line THP-1 (TIB-202, ATCC) was cultured in RPMI-1640 medium (Gibco) supplemented with 10 % FBS and 4 mM L-glutamine at 37°C in a humidified 5 % CO_2_ atmosphere. ATCC routinely performs cell line authentication using short tandem repeat profiling. Cell experimentation was always performed within a period not exceeding 20 passages after resuscitation and cells were regularly tested for mycoplasma.

To differentiate THP-1 cells into macrophages, we cultured THP-1 cells at a concentration of 5×10^5^ cells/mL with 10 ng/mL Phorbol 12-myristate 13-acetate (PMA) (Sigma-Aldrich) for 48 h, followed by 24 h with fresh medium without PMA.

These culture conditions were optimised by evaluating cell viability and immunophenotype using CD11b, CD11c, CD14, and CD44 (**Supplementary Fig. 1**). We assessed cell viability by propidium iodide staining.

### Bacterial strains and culture conditions

*Salmonella enterica* serovar Typhimurium SL1344 and derived Δ*phoP* mutant (**Supplementary Table 1**) were kindly provided by Dirk Bumann (Biozentrum Basel) and grown in LB broth at 37°C with constant rotation. All other mutants were constructed by gene deletion using the λ red recombinase system^51^ with modifications^52^ using the primers described in **Supplementary Table 2**.

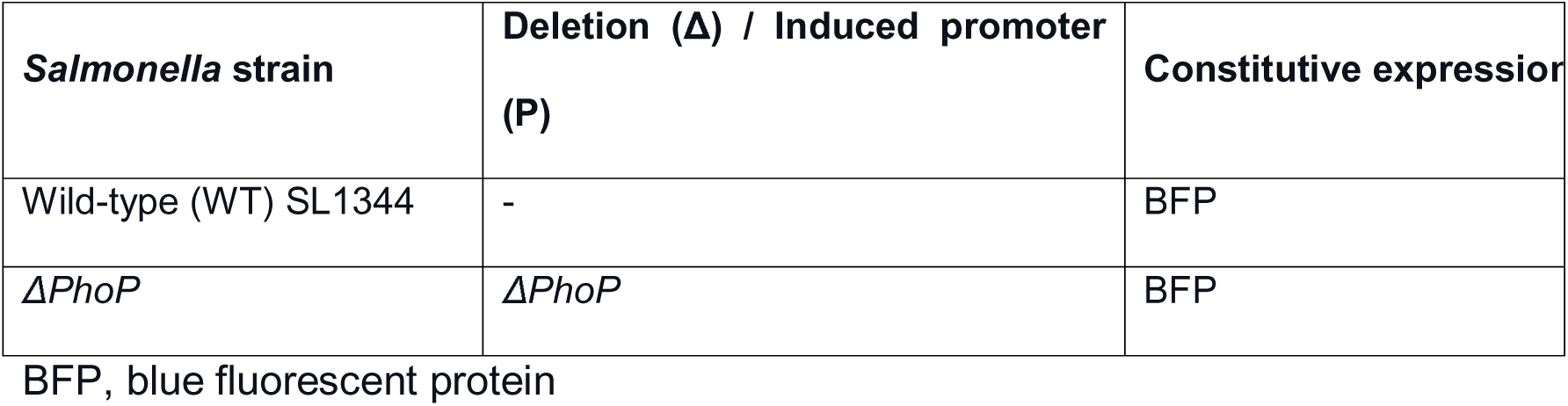

| ***Salmonella* strain** | **Deletion ($\Delta$) / Induced promoter (P)** | **Constitutive expression** |
| --- | --- | --- |
| Wild-type (WT) SL1344 | - | BFP |
| $\Delta PhoP$ | $\Delta PhoP$ | BFP |

BFP, blue fluorescent protein

**Table 2.**
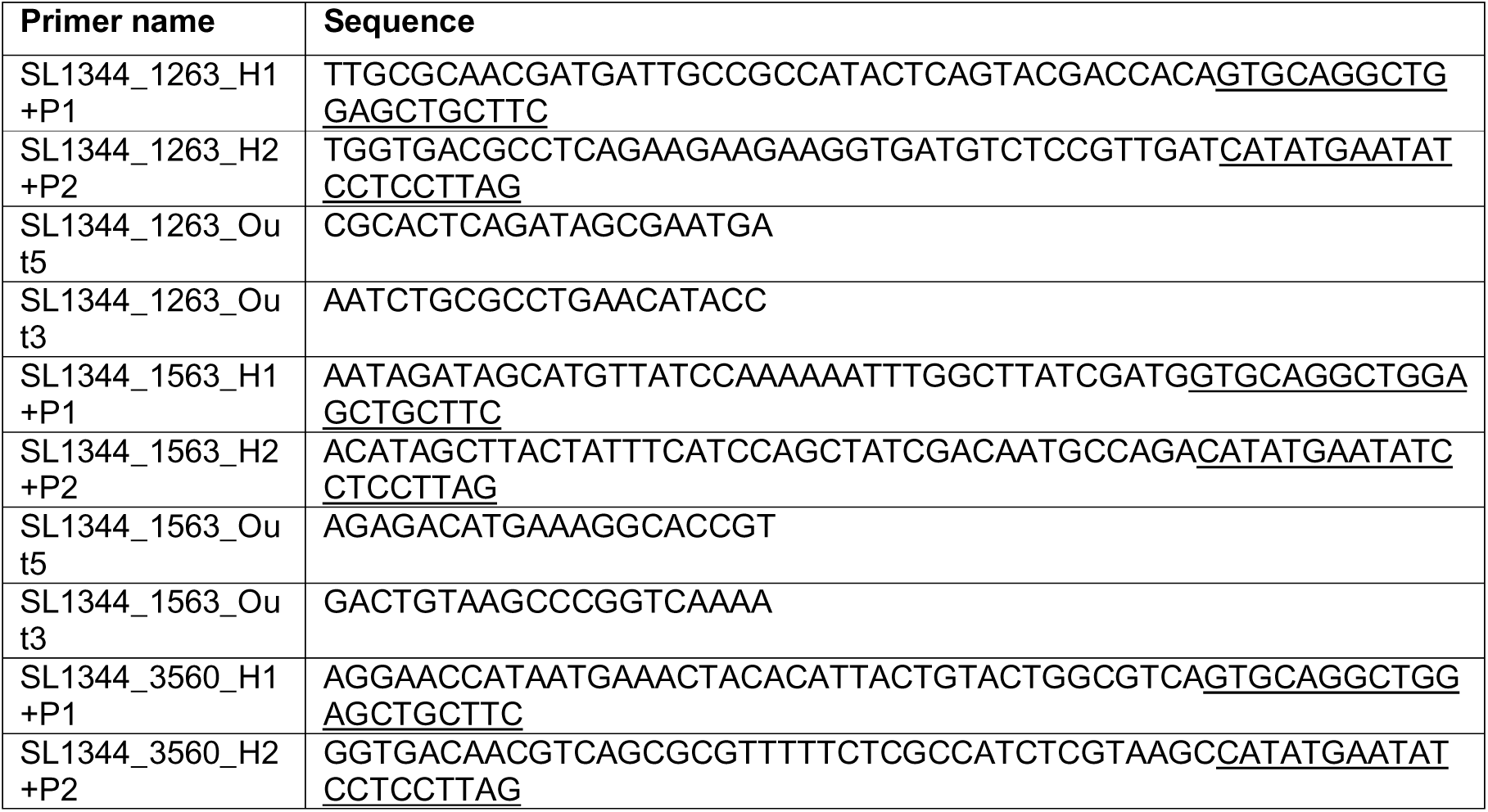
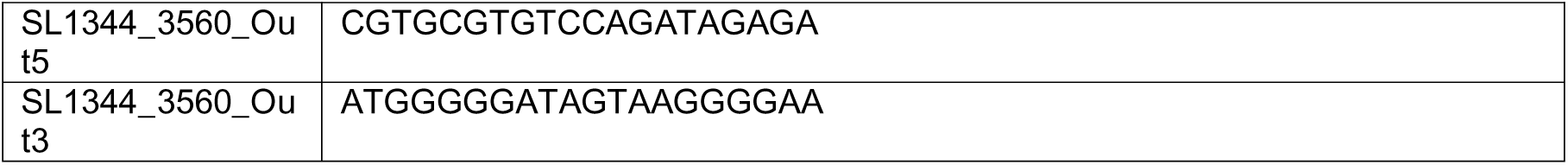
Primers used in this study.

Underlined sequences correspond to the region that anneals to the 5′ or 3′ end of the kanamycin-resistance cassette in the template vector pCLF4 (GeneBank accession number HM047089).

### Bacterial infection

For each experimental replicate, 4×10^7^ THP-1 cells were differentiated into macrophages, seeding 1×10^7^ cells per 150 mm tissue culture dish. Before infection, the cell medium was refreshed with 10 mL of complete medium (washed three times with warm PBS before refreshing the medium). Overnight bacteria cultures (optical density (OD600 nm) of 1.0 – 1.2, where OD= 1 is equals to approximately 1×10^9^ bacteria/mL), were washed in warm PBS twice at 4,000 rpm for 15 min and, subsequently, resuspended in warm cell culture medium at the desired MOI of 100 at a concentration of 1×10^9^/mL. Bacterial input was placed on LB agar plates to count colony-forming units (CFUs). Then bacteria were added at an MOI of 1:100 and incubated for 30 min for uptake. This produced the 30 min time-point. After 30 min, we removed the bacteria-containing medium, washed twice with warm PBS and refreshed the medium with 50 μg/mL gentamicin. After one hour, cells were rewashed twice with warm PBS and incubated with fresh medium supplemented with 15 μg/mL gentamicin up to 4 h (4 h post-infection or p.i.). CFUs were determined by lysing cells in 0.1 % Triton-X 100 in PBS at indicated time points.

### Bacterial staining with CellTrace Violet (CTV)

Before the infection, untagged bacteria were stained with different concentrations of CellTrace Violet (CTV) dye (Thermo-Fisher Scientific), following the manufacturer’s protocol. Higher concentration of CTV dye tested, 500 μM, stained 97 % of bacteria, and its fluorescence intensity was compatible with mTagBFP. Moreover, CTV dye did not affect bacteria uptake (**Supplementary Fig. 2B**). Briefly, the OD_600_ was measured for overnight culture and about 10^9^ bacteria were incubated in 500 μl of warm PBS with 500 μM of CTV stain (resuspend immediately to achieve homogenous staining) for 20 min at 37°C. Then, the bacteria were washed twice with PBS (at 37°C) for 5 min and centrifugated at 4,000 rpm for 15 min. Finally, bacteria were resuspended in a warm cell culture medium at a MOI of 100.

### Cell lysis for phagosome isolation

After the indicated time point (30 minutes for uptake and 4 h p.i.), the medium was removed, and ice-cold PBS was added. Then, cells were scraped from four dishes, transferred into 15 mL tubes, and centrifuged at 300 xg at 4°C for 10 min. Cell pellets were washed twice with ice-cold PBS. Afterwards, pellets were re-suspended in 1 mL hypotonic buffer (250 mM sucrose, 3 mM imidazole, pH 7.4) with protease and phosphatase inhibitors and 0.1µl (25 U) DNase (Pierce Universal Nuclease for Cell Lysis, Thermo-Fisher Scientific). To lyse the cells, we used a Dounce homogeniser in a class II biological safety cabinet (between 5 – 7 strokes until about 70% of cells were broken). We monitored the lysis using trypan blue dye under the microscope. Cell lysates were centrifuged at 1,200 xg at 4°C for 10 min to pellet nuclei. The post-nuclear supernatant I was then transferred into a 2 mL tube, and again treated with DNase for 5mins at 37°C and the nuclei were pelleted again at 1,200 x g for 5 mins. The post-nuclear supernatant II was then transferred into a fresh 2 mL tube and diluted with the same volume of ice-cold PBS and centrifuged at 2,500 xg at 4°C for 5 min to pellet phagosomes.

### CellVue Claret Far Red staining

We used the CellVue Claret Far Red Membrane Label (Sigma-Aldrich) for phagosomal membrane staining as per manufacturer’s protocol. In brief, we resuspended the phagosome pellet in 50 μl of diluent C. Then, 50 μl with 4 μM dye was added, mixed and incubated for 5 min at room temperature. The staining was quenched using 100 μL of 1% BSA in PBS and washed twice with ice-cold PBS by centrifugation at 2,500 xg at 4°C for 5 min. Samples were fixed with 1% PFA in PBS at 4°C for 15 min, followed by one washed with PBS. Finally, we resuspended the samples in 1 mL of PBS with 1 mM EDTA for flow cytometry.

### Flow cytometry

For immunophenotype analysis, cells were blocked with flow cytometry staining buffer at 4°C for 20 min, then incubated with the corresponding antibody at a 1:100 dilution at 4°C for 20 min, washed with PBS, and analysed using a BD Symphony A5 flow cytometer (BD Biosciences).

The samples were then flow sorted for double positives (BFP+ and Claret Far Red+) on a BD FACS Aria III. The BFP excitation was set to 399 nm and emission to 454 nm and for Cell Vue Claret Far Red excitation to 655 nm and emission to 675 nm. The initial gating strategy involved the selection for bacterial phagosome size followed by gating based on single channel controls. All samples were sorted based on a double positive gating channel and the analysis was performed using FlowJo software.

### Sample preparation for LC-MS/MS

After sorting, phagosomes were pelleted by centrifugation at 16,000 xg for 15 min at 4°C, resuspended in PBS supplemented with protease inhibitors, transferred to low protein binding microcentrifuge tubes. Next, the samples were resuspended in a 50 μL lysis buffer (5% SDS and 50 mM triethylammonium bicarbonate (TEAB) in HPLC water, pH 7.4) supplemented with protease inhibitors (cOmplete, EDTA-free, Roche). To prepare the samples for mass spectrometry, S-Trap micro columns (Protifi) were used according to the manufacturer’s recommended protocol for high recovery. Proteins were reduced with 20 mM tris(2-carboxyethyl)phosphine at 37°C for 30 min and alkylated with 20 mM iodoacetamide in the dark for 30 min. The proteins were then acidified with 5 μL 55 % phosphoric acid and 330 μL S-Trap binding buffer. 1 microgram of Sequencing Grade Modified Trypsin (Promega) were added to the acidified samples before being transferred into the S-Trap column. The columns were washed five times with 150 μL S-Trap binding buffer before incubating them with 0.5 μg of Sequencing Grade Modified Trypsin in 25 μl of 50 mM TEAB, pH 8 for 2 h at 47°C. Finally, the samples were sequentially eluted in 40 μl 50 mM TEAB, pH 8, 40 μl 0.2 % formic acid and 35 μl 50 % acetonitrile with 0.2 % formic acid. The eluted peptides were dried in a vacuum concentrator and stored in −80°C until further use.

### Data-independent acquisition mass spectrometry (DIA-MS)

Liquid chromatography (LC) was performed using an Evosep One system with a 15 cm Aurora Elite C18 column with integrated captive spray emitter (IonOpticks), at 50 °C. Buffer A was 0.1 % formic acid in HPLC water, buffer B was 0.1 % formic acid in acetonitrile. Immediately prior to LC-MS, peptides were resuspended in 30 µl of buffer A and a volume of peptides equivalent to 20 % of the sample (100,000 events isolated during the FACS) was loaded onto the LC system-specific C18 EvoTips, according to manufacturer instructions, and subjected to the predefined Whisper100 or Whisper-Zoom 20 SPD protocol (where the gradient is 0-35 % buffer B, 100 nl/min or 200 nl/min, for 58 minutes). The Evosep One (EvoSep) was used in line with a timsTOF-HT mass spectrometer (Bruker) operated in DIA-PASEF mode with high sensitivity on. Mass and IM ranges were 300-1200 *m/z* and 0.6-1.45 1/*K_0_*, DIA-PASEF was performed using variable width IM-*m/z* windows without overlap, designed using py_diAID^11^, and are provided (see data sharing section). TIMS ramp and accumulation times were 100 ms, total cycle time was ∼1.8 seconds. Collision energy was applied in a linear fashion, where ion mobility = 0.6-1.6 1/K_0_, and collision energy = 20 - 59 eV.

### MS data analysis

Raw data files were searched using DIA-NN 1.9,^53^ using its *in silico* generated spectral library from the *Homo sapiens* (downloaded from Uniprot on the 14th of September 2023, containing 42437 entries) and *Salmonella* Typhimurium (downloaded from Uniprot on the 14th of September 2023 containing 4659 entries) proteomes and a common contaminants list^54^. Carbamidomethylation (C) was set as a fixed modification, and oxidation (M) to variable. Precursor masses from 300 to 1800 were included, with charge 1 to 4. Mass accuracy for MS1 and MS2 was set to 15, Match between runs was performed and normalisation set to ‘Global’. Trypsin specificity was set to one missed cleavage and a protein and PSM false discovery rate of <1%, respectively. Stringent filtering was applied: Proteins required intensities in a minimum of 3/6 in one group, and 0,3-6 in the other and proteins required to have a minimum of 2 peptides.

Proteomics data analysis was performed on a R software pipeline (v4.2.1). Raw intensities were log2-transformed. Protein groups detected in at least four samples within any experimental group were kept for downstream analysis. For some data, missing values were imputed by v2-mnar method implemented in msImpute package^55^ (v1.7.0). Differential expression between groups was assessed by fitting a linear model and applying empirical Bayes moderated t-statistics in limma package^56^ (v3.60.4). Resulting P-values were adjusted using the Benjamini–Hochberg FDR correction. Proteins with a log2-fold change ≥ ±0.5 and adjusted p-value ≤ 0.05 were considered as differentially expressed.

### Rotenone treatment of infected macrophages

Murine bone marrow derived macrophages (BMDM) were isolated from at least four C57BL/6J wild type mice per experiment using L929 supplemented medium ^57,58^. BMDM were seeded with the density of 3X10^5^ per well of a 24 well plate

Wild-type (WT) *Salmonella enterica* serovar Typhimurium (STM) SL1344 was cultured overnight in LB supplemented with 100 mg/ml kanamycin at 37C and the OD_600_ was used to calculate the bacteria concentration. The bacteria were washed twice with PBS and then resuspended in pre-warmed media and BMDM infected with WT STM at multiplicity of infection MOI 20:1 as described above, however, 2 µM rotenone or vehicle (PBS) was added with the gentamicin step. The STM-infected macrophages were lysed using 1% triton X-100 and the colony formation units (CFU) determined on kanamycin agar plates overnight.

### Data availability statement

The mass spectrometry proteomics data have been deposited to the ProteomeXchange Consortium via the PRIDE partner repository^59^ with the data set identifier PXD062621.

## Acknowledgements

This research was funded by a Wellcome Trust Investigator Award (215542/Z/19/Z) to MT, a Marie Skłodowska-Curie and a Swiss National Science Foundation Fellowship to FRC, and a BBSRC studentship to MM. The mass spectrometer used in this work was funded by a Wellcome Trust multi-user equipment grant (212947/Z/18/Z). J.E. and C.V. acknowledge funding from the ANR (Agence Nationale pour la Recherche) through the grant “PureMagRupture”. The Enninga team is part of the LabExes “IBEID” and “Milieu Interieur”. We thank the Flow Cytometry Core Facility (FCCF) of Newcastle University for their support.

## Notes

### Competing Interest Statement

The authors have declared no competing interest.

### Summary of Updates

Updated manuscript, removal of one dataset (dead STM) for simplicity. some new experiments with mutants and mitochondrial inhibitor.

