## Supplementary Figures 1-5 for "Flow cytometry-based isolation of *Salmonella*-containing phagosomes combined with ultra-sensitive proteomics reveals novel insights into host-pathogen interactions"

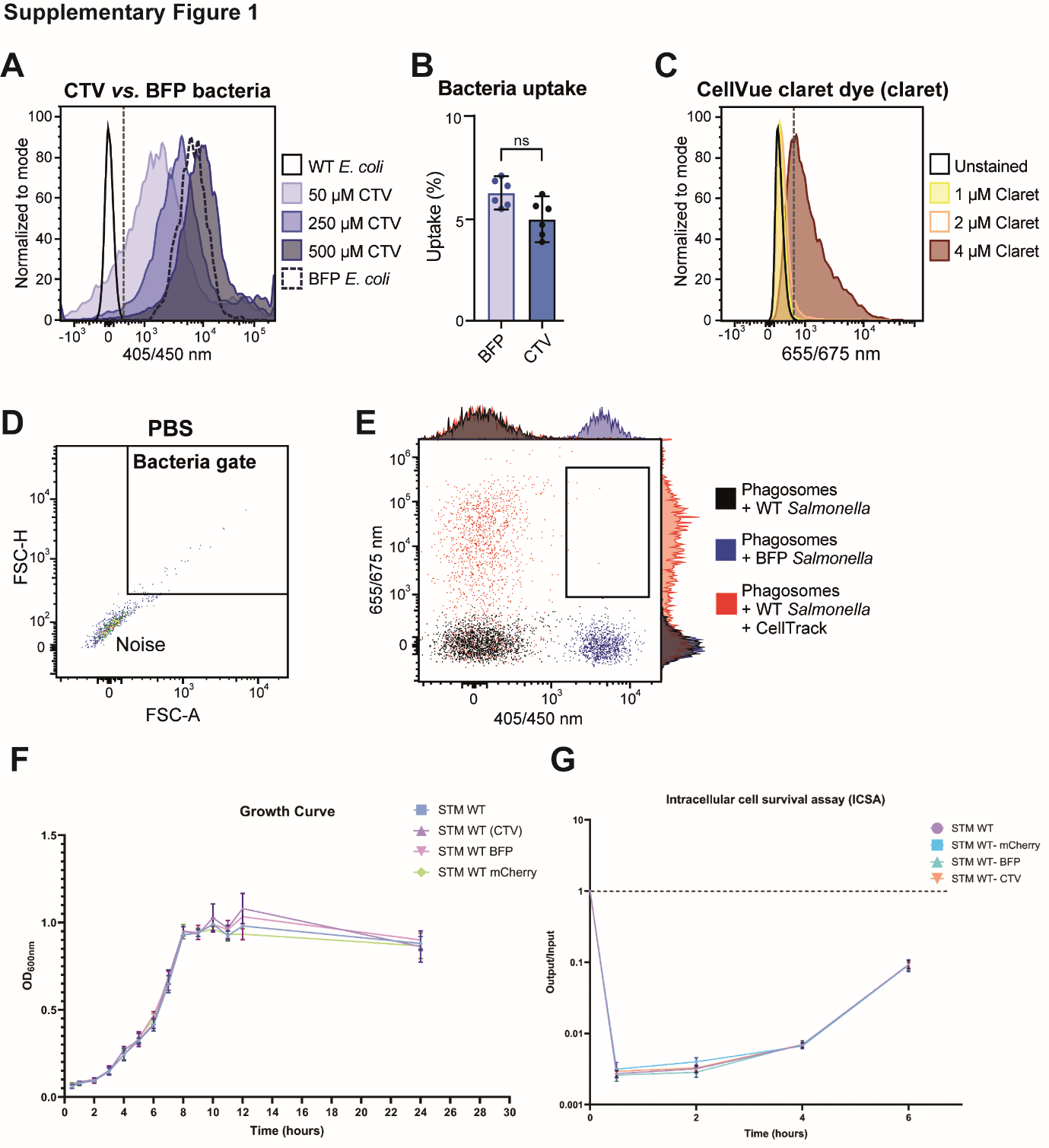

**Supplementary Figure 1: Optimisation of phagosome staining protocol. A)** Unstained wild-type (WT) DH5α *E. coli* were stained with 50, 250 and 500 μM CellTrace Violet (CTV) and compared with mTagBFP *E. coli* (BFP) (Ex/Em 405/450 nm). **B)** Staining with CTV does not affect phagocytic uptake. **C)** Phagosomes from THP-1 cells were stained with 0 (unstained), 1, 2 and 4 μM CellVue Claret Far Red dye (claret, Ex/Em 655/675 nm). THP-1 monocytes were treated with 10, 50, or 100 ng/mL PMA for 48 h following for 24h with fresh medium without PMA (orange bars) or for 72 h of continuous treatment with PMA (blue bars). **D)** PBS was used to determine the electronic noise of the cytometer and sorter. **E)** Phagosomes with WT (i.e. non-fluorescent) Salmonella (black), with BFP Salmonella (blue) (but no CellVue staining), and with WT Salmonella and CellVue claret (red) were analysed as controls to determine the gate of double positive events (claret and BFP).

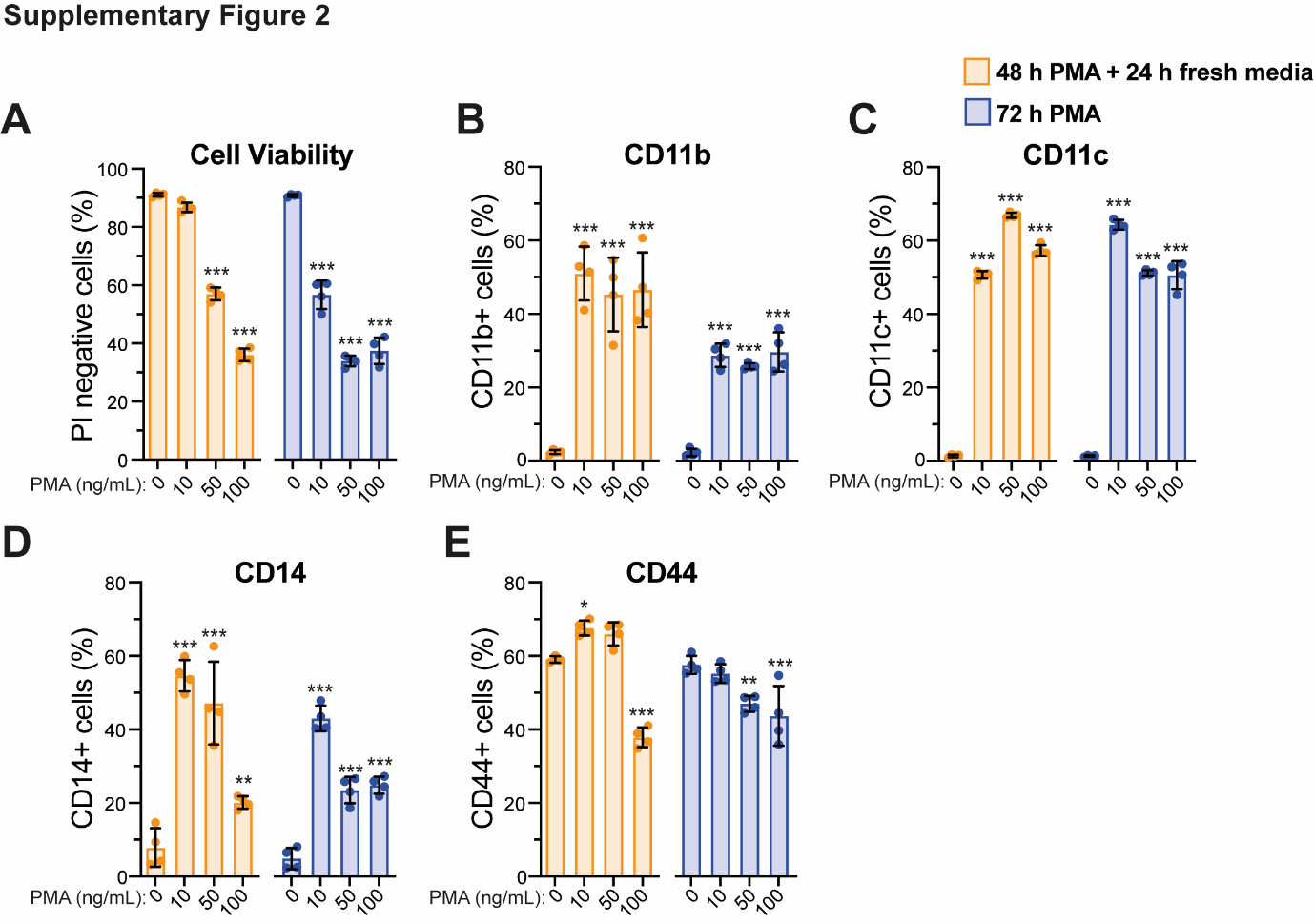

**Supplementary Figure 2: Differentiation of THP-1 monocytes into macrophages**

THP-1 monocytes were treated with 10, 50, or 100 ng/mL PMA for 48 h following for 24h with fresh medium without PMA (orange bars) or for 72 h of continuous treatment with PMA (blue bars). **A)** Percentage (%) of negative cells for propidium iodide (PI). Percentage of cells positive for **B)** CD11b, **C)** CD11c, **D)** CD4, **E)** CD44 measured by flow cytometry. Two-way ANOVA test followed by Dunnett's multiple comparisons test was used for the comparisons with untreated samples. The statistical significance is indicated as follows: ***, p <0.001; **, p<0.01; *, p<0.05. Error bars represent the standard deviation of four independent replicates.

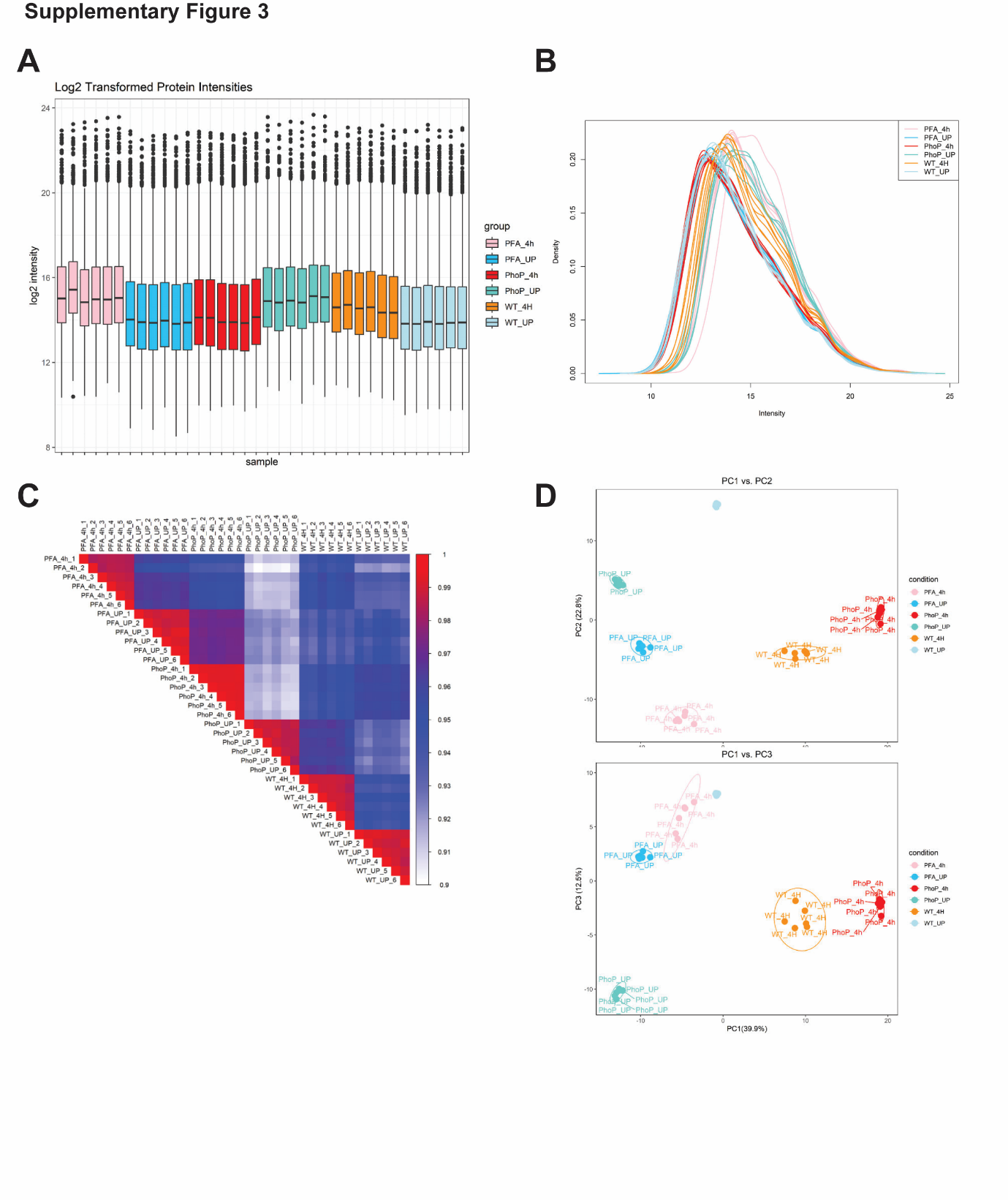

**Supplementary Figure 3:** **PhagoCyt proteomics data from *Salmonella* infected THP-1 cells phagosomes**. **A)** Box and whisker plot of the log2 transformed normalised LFQ intensities of six biological replicates per group (STM WT uptake (30 min), STM WT 4h p.i., STM *ΔphoP* uptake, STM *ΔphoP* 4h p.i., STM WT PFA Fixed (dead) uptake and STM WT PFA Fixed (dead) 4h), **B)** Kernel density plot, **C)** Pearson‘s correlation plot and **D)** Principal component analysis (PCA) of the whole proteome dataset shows the three main principal components (PC).

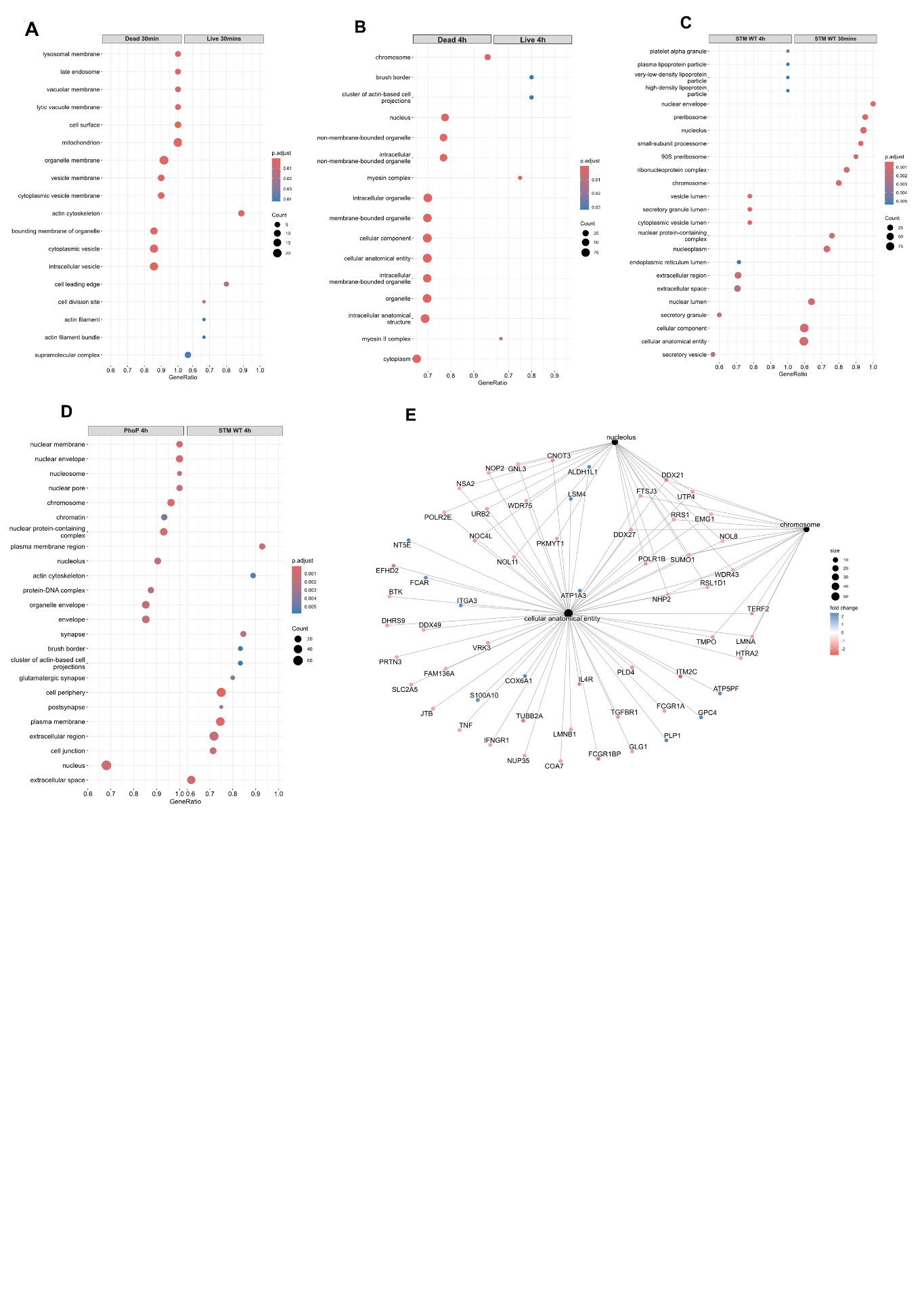

**Supplementary Figure 4:** **Gene set enrichment analysis (GSEA) of the PhagoCyt proteomics data from *Salmonella* infected THP-1 cells phagosomes**. **A)** Gene ontology of the cellular compartment analysis using R-studio cluster profiler package for top 100 genes enriched in **A)** STM PFA fixed (dead) vs STM WT (live) at 30mins p.i.; **B)** STM PFA fixed (dead) vs STM WT (live) at 4 h p.i.; **C)** STM WT 4h vs 30mins p.i.; **D)** STM WT vs STM *ΔphoP* at 4 h p.i.; **E)** CNET plot for STM WT 4h p.i. phagosomes for top 100 genes using R-studio cluster profiler package. *Guangchuang Yu, Li-Gen Wang, Guang-Rong Yan, Qing-Yu He. DOSE: an R/Bioconductor package for Disease Ontology Semantic and Enrichment analysis. Bioinformatics 2015, 31(4):608-609*

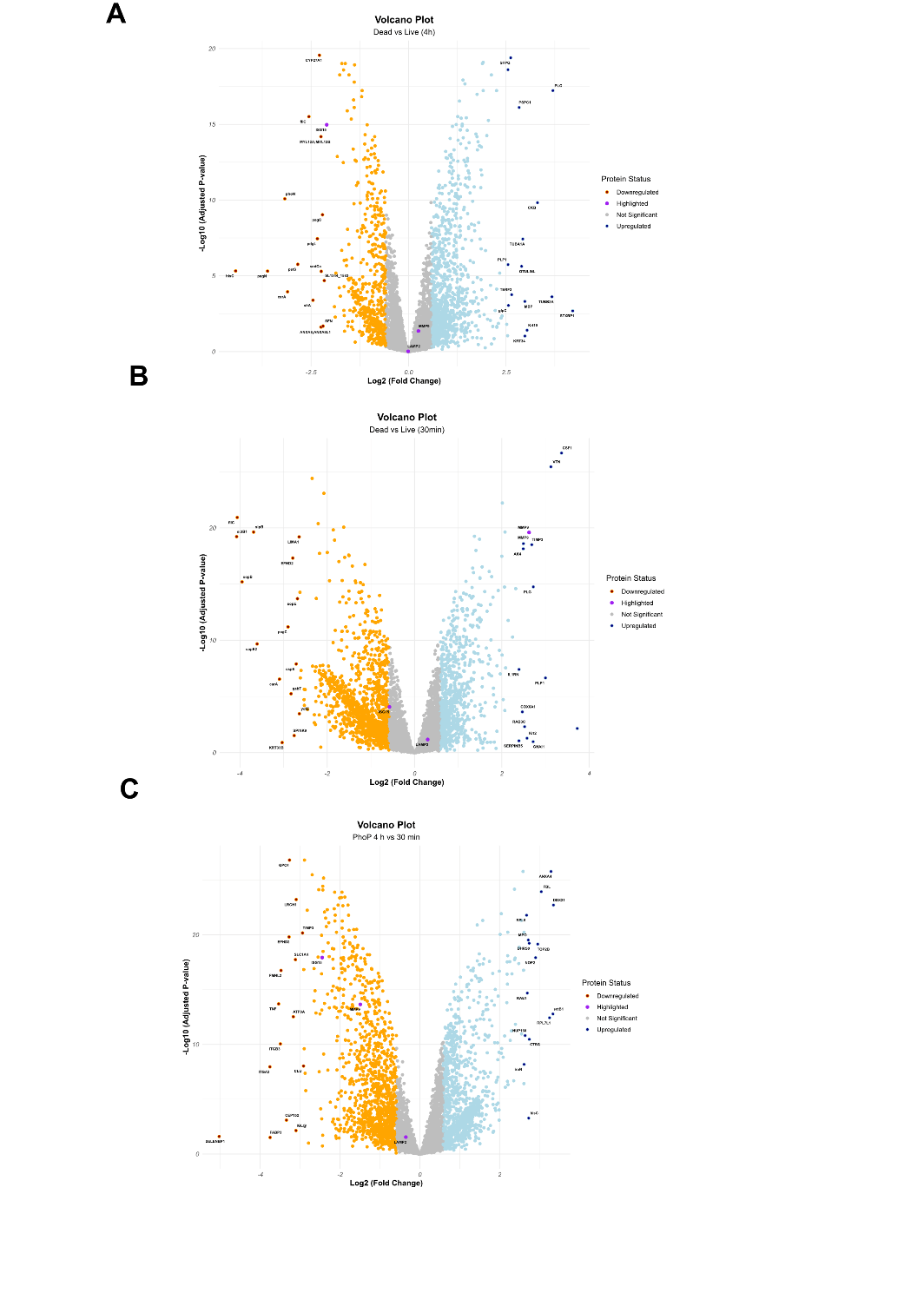

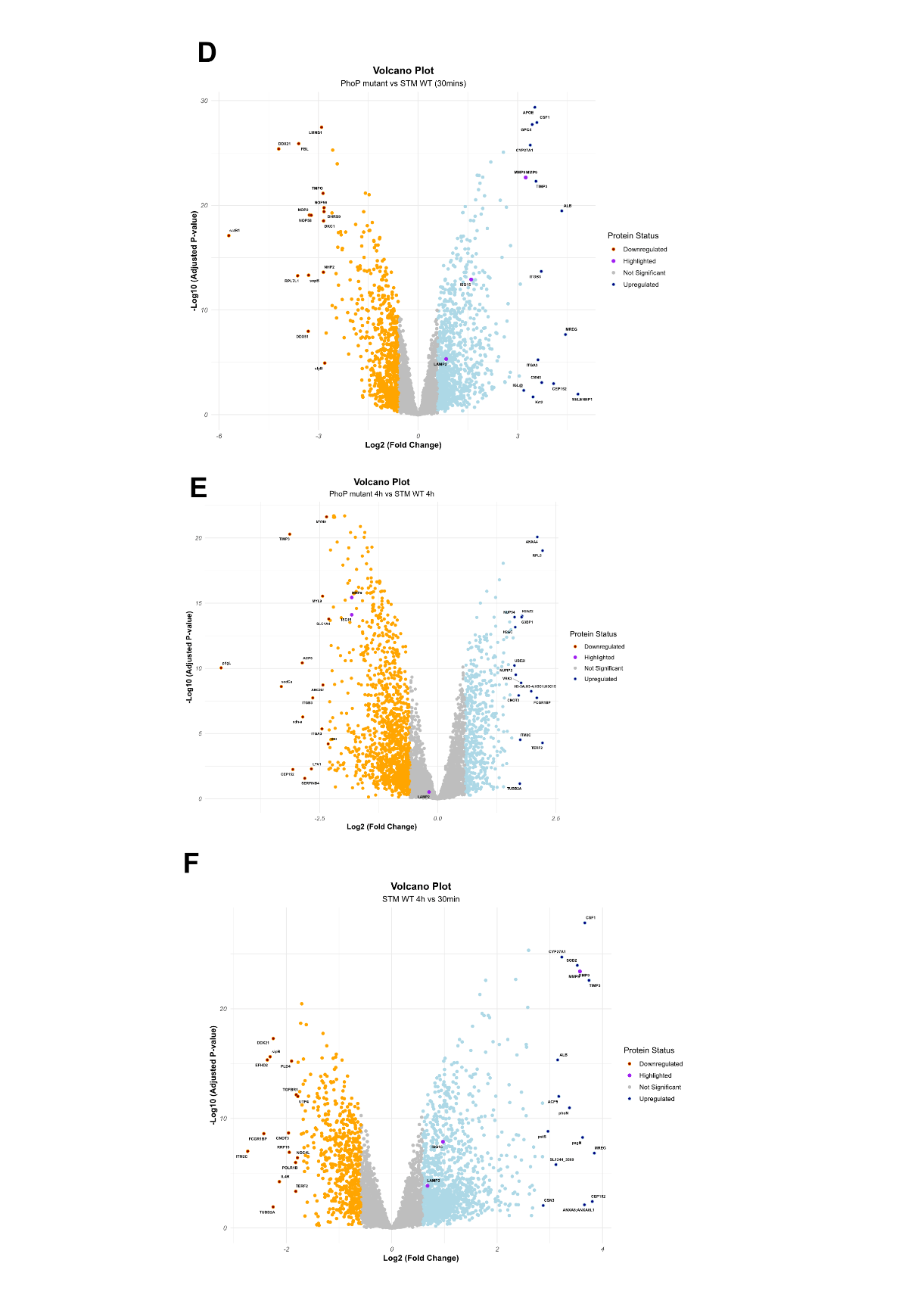

**Supplementary Figure 5: Volcano plots of phagosome proteome comparisons of WT, ΔphoP and dead STM.**

**A)** The volcano plot displays the downregulated proteins (in orange) and upregulated proteins (in blue) that are significant in WT *Salmonella* 4h p.i compared to uptake phagosomes in **A**) Dead vs live at 4 h, **B**) Dead vs live 4h , **C**) PhoP mutant 4h vs 30mins , **D**) PhoP 4h vs WT 4h, **E**) PhoP vs WT 4h, and **F**) WT 4h vs WT 30mins. Proteomics dataset is analysed by limma and graphs are plotted using R-studio.
